# Correlative Imaging of 3D Cell Culture on Opaque Bioscaffolds for Tissue Engineering Applications

**DOI:** 10.1101/2023.03.17.533202

**Authors:** Monet Sawyer, Joshua Eixenberger, Olivia Nielsen, Jacob Manzi, Raquel Montenegro-Brown, Harish Subbaraman, David Estrada

## Abstract

Three-dimensional (3D) tissue engineering (TE) is a prospective treatment that can be used to restore or replace damaged musculoskeletal tissues such as articular cartilage. However, current challenges in TE include identifying materials that are biocompatible and have properties that closely match the mechanical properties and cellular environment of the target tissue, while allowing for 3D tomography of porous scaffolds as well as their cell growth and proliferation characterization. This is particularly challenging for opaque scaffolds. Here we use graphene foam (GF) as a 3D porous biocompatible substrate which is scalable, reproduceable, and a suitable environment for ATDC5 cell growth and chondrogenic differentiation. ATDC5 cells are cultured, maintained, and stained with a combination of fluorophores and gold nanoparticle to enable correlative microscopic characterization techniques, which elucidate the effect of GF properties on cell behavior in a three-dimensional environment. Most importantly, our staining protocols allows for direct imaging of cell growth and proliferation on opaque GF scaffolds using X-ray MicroCT, including imaging growth of cells within the hollow GF branches which is not possible with standard fluorescence and electron microscopy techniques.

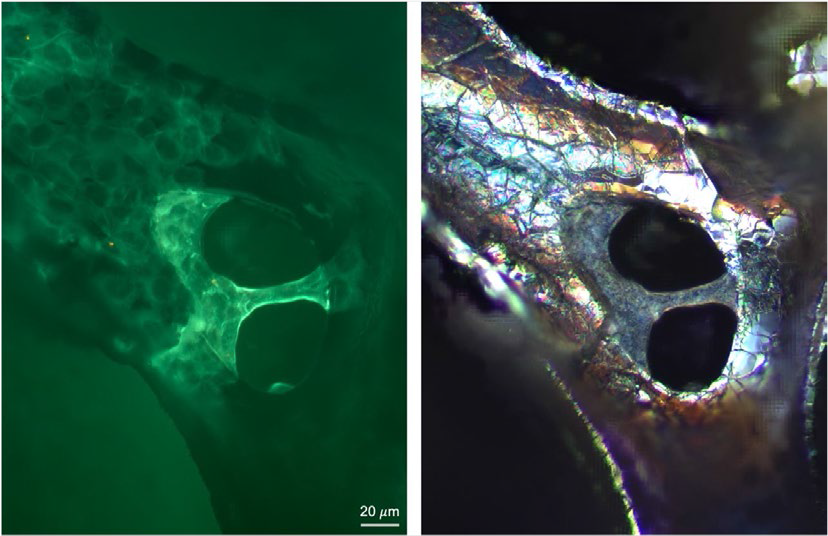

---

Articular cartilage (AC) damage is a frequent occurrence that can lead to osteoarthritis, the most prevalent joint disease and leading cause of disability in the United States and other developed nations. Tissue engineering (TE), a prospective alternative treatment for this musculoskeletal disorder, aims to repair, maintain, or regenerate these damaged tissues; however, human tissues are complex in function, structural hierarchy, and scale, making it difficult to synthesize functional tissue in a lab that can be used for clinical treatments^1^. Although there have been advances in engineering AC, significant barriers remain regarding the ability to generate functional articular cartilage that imitates native cartilage both in structure and mechanical function.

Over the last decade, AC TE has evolved from two-dimensional cell cultures grown on planar surfaces to culturing cells in complex three-dimensional architectures^2^. The goal with next-generation bioscaffolds is to closely mimic the native environment of AC as it is known that the physical properties of bioscaffolds can drive specific cell behaviors such as proliferation, differentiation, and extracellular matrix (ECM) production leading to tissue formation^3–5^. In addition, bioscaffolds that have materials properties that can be used to drive cell behavior by generating physical cues to stimulate tissue growth are advantageous for AC TE^6^.

Graphene foam (GF) is a 3D porous biocompatible substrate that is easily produced via chemical vapor deposition (CVD) on a nickel template. It’s unique material properties such as high electron mobility, excellent thermal conductivity, and high mechanical strength, can be taken advantage of to drive cell behavior, while synthesis via the CVD process lends to its scalability^7,8^. Further, the microporous structure and high porosity of GF facilitates nutrient exchange, and the high surface/volume ratio provides a favorable environment for long-term cell attachment and growth, making it an optimal candidate for a bioscaffold that meets the demands of new-generation three-dimensional biomaterials^9^.

Although 3D environments are more conducive to functional tissue formation, characterization of cell proliferation and migration in these systems remains a challenge^10^. Unlike 2D cell cultures, analyzing a single cell plane is not sufficient when working with three-dimensional systems as it important to assess proliferation and migration of cells within the bioscaffold as it relates to bioscaffold porosity, structure thickness and pore interconnectivity. A high cell density and even spatial distribution is associated with functional tissue formation; therefore, it is important to accurately evaluate cell attachment as well as cell distribution and density after seeding^5^. Electron and confocal fluorescence microscopy common methods used to analyze bioscaffold structure and cell behavior within the scaffold, however, these methods do not allow for a three-dimensional analysis of the bioscaffold environment corresponding to its internal structure and are limited by the bioscaffold’s opacity. Microcomputed tomography (MicroCT) techniques have been developed to study bioscaffold architecture without sample damage^10^, however, the investigation of cellular activity in a 3D environment is difficult due to their low contrast as compared to scaffold materials. Here we have demonstrated a method for culturing, maintaining, and labelling cells grown on GF bioscaffolds to quantify the effect of GF properties on spatial cellular distribution in a three-dimensional environment through MicroCT analysis.

We utilize our open-source CVD furnace to synthesize GF bioscaffolds using a nickel foam template^11^. Following synthesis, the Gr/Nickel foam substrates are post-processed in 3 M hydrochloric acid until complete dissociation of the nickel. To characterize the superficial microstructure, we imaged the GF using scanning electron microscopy (FEI Teneo Field Emission Scanning Electron Microscope) which shows the macroporous structure and wrinkled topography with increasing magnification and microcracks in the branches due to the dissociation and internal removal of the nickel foam template indicated in Figure 1B. GF was further analyzed with X-ray Raman spectroscopy which verified complete dissociation of the nickel foam template and the graphitic nature of our CVD GF (Figure 1C). Raman spectra was compared at three separate locations across a single sample, further confirming that the quality of GF is consistent throughout the bulk.

**Figure 1.**
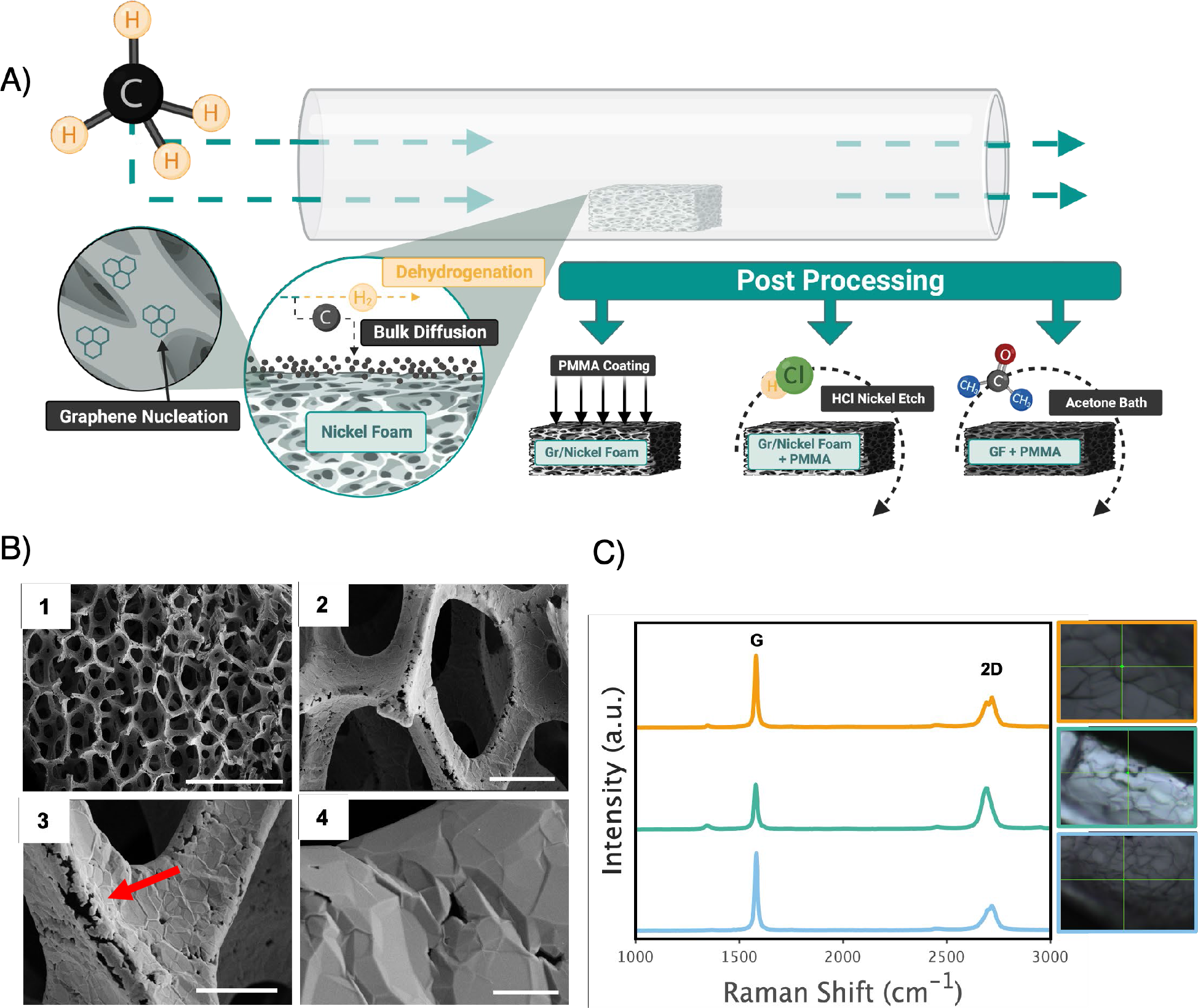
GF synthesis and characterization. A) GF is synthesized via CVD on a nickel foam template. Gr/Nickel foam is coated in PMMA to maintain integrity during 3M HCl bath to dissolve nickel template. After nickel dissociation, PMMA is disolved in an acetone bath. (B) SEM micrographs showing the bulk structure and wrinkled topography of CVD GF with scale bars of (1) 1 mm, (2) 100 μm, (3) 50 μm, and (4) 5 μm, a red arrow indiating micrcracks in the branch. (C) Raman spectroscopy indicates graphitic quality with the characteristic G and 2D peaks seen in graphitic materials as well as complete nickel dissociation.

To prepare our GF bioscaffolds for ATDC5 (Sigma Aldrich) cell culture, we sterilized them with ethanol prior to conditioning them in growth media composed of F12/Dulbecco’s Modified Eagle Medium (DMEM:F12), 5% (v/v) fetal bovine serum (FBS), and 1% (v/v) penicillin/streptomycin for 24 hours before seeding them with ATDC5 cells. We used an anti-adherence rinsing solution (STEMCELL technologies) in our well plates to prevent cell growth on the culture ware containing the GF and promote cell growth on the scaffold. Conditioned GF bioscaffolds were seeded with 5×10^6^ ATDC5 cells and incubated for 7-days in growth media at 37°C and 5% CO_2_.

Cells were fixed on GF bioscaffolds with 0.2% PFA, permeabilized with 0.1% Triton-X, and indirectly labelled with beta-Actin polyclonal antibody (Thermo Fisher Scientific) and Goat anti-Rabbit IgG (H+L) Secondary Antibody Alexa Fluor™ 488-10 nm colloidal gold (Figure 2A). After an incubation period, bioscaffolds were rinsed 10 times with phosphate buffered saline (PBS) diluted in nanopure water (10:1). The secondary antibody was vortex mixed to ensure even dispersion of gold nanoparticles before it was added to GF bioscaffolds, incubated, and rinsed 10 times with the same PBS dilution. After labelling, we sought to assess the success of our labelling procedure through transmitted light and fluorescence microscopy to confirm that the fluorophore had stained the actin of the cell. In Figure 2C and D we utilized Circular polarized light - Differential Interference Contrast (C-DIC), a reflected light technique, which converts gradients in the specimen optical path into sample amplitude differences and allows for the visualization of the wrinkled topography of GF bioscaffolds which were then colocalized with fluorescence micrographs as seen in Figure 2F and G. Cells spanning GF pores are optically dense enough to see using transmitted light microscopy techniques and actin labelling allows for the quantification of cell attachment on the GF surface, however, GF is optically opaque (Figure 2B) making it impossible to quantify cell migration and dispersal throughout the scaffold using traditional methods such as confocal fluorescence microscopy. Scanning electron micrographs were taken after indirect labelling, as the final confirmation of our straining technique, and allowed for the quantification of cell attachment and morphology on GF with SEM and no additional processing of the samples, affirming that the gold nanoparticles were conjugated to the antibody that we used to label the actin of the cell. After validating our staining technique, we performed MicroCT (SkyScan 1172 Xray MicroCT) on our labelled samples to determine ATDC5 distribution and migration within the three-dimensional environment.

**Figure 2.**
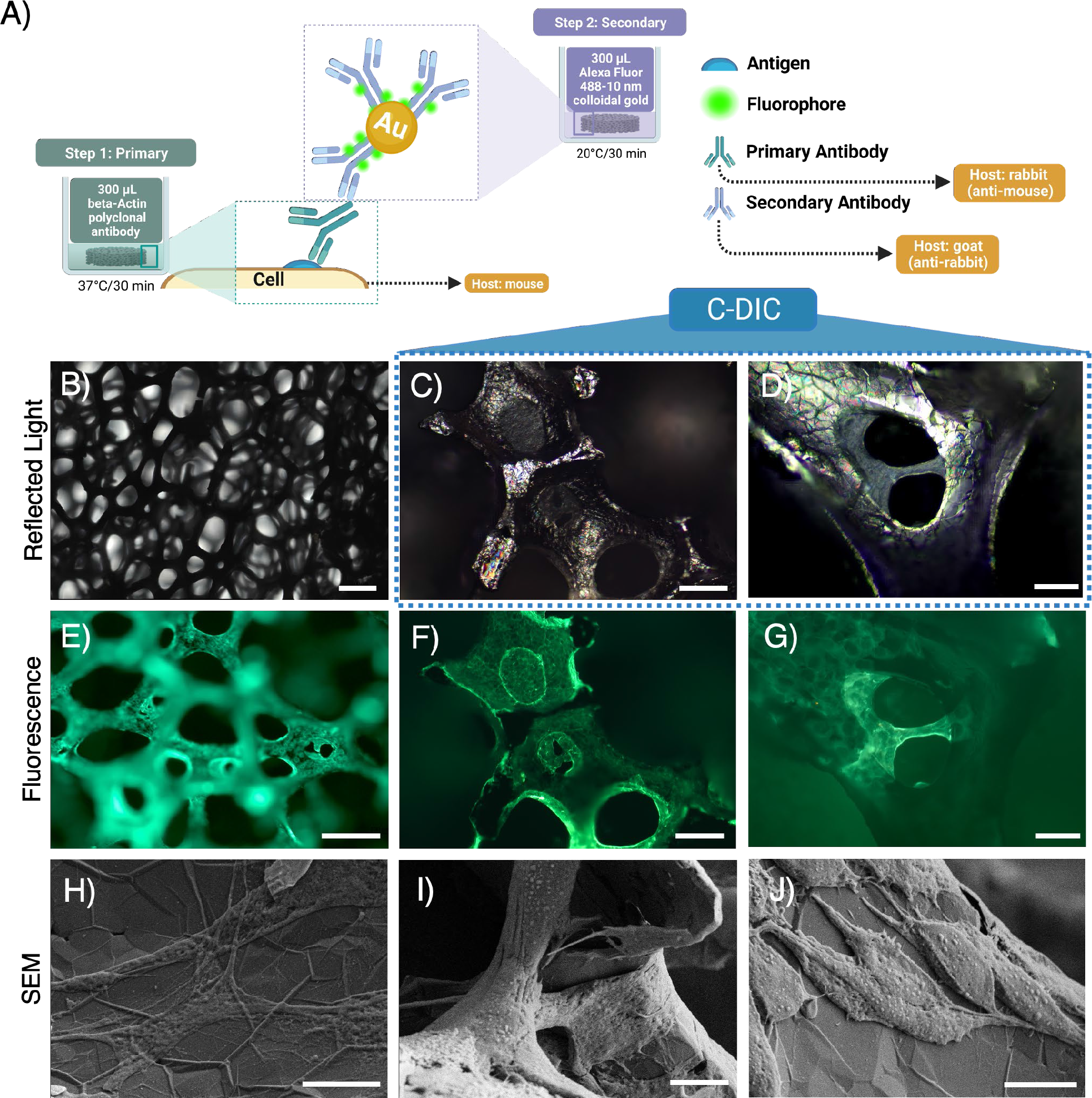
Indirect labelling of ATDC5 cells on GF bioscaffolds. A) Labelling procedure and incubation parameters for double labeling with beta-Actin polyclonal antibody and Goat anti-Rabbit IgG (H+L) Secondary Antibody Alexa Fluor™ 488-10 nm colloidal gold. B) Reflected light micrographs exhibit GF’s opaque quality with limited z-resolution. C-D) C-DIC shows the wrinkled structure of GF and ATDC5 cells spanning pores colocalized with fluorescence images (F-G). H-J) SEMs confirm that the gold nanoparticles were conjugated with the secondary antibody allowing for the visualization of cell attachment and morphology.

MicroCT acquisition on bare and labelled GF bioscaffolds was conducted with a source voltage, current, and pixel size of 26kV, 146 μA, and 2.24 μm respectively. The exposure time was set to get quality images with a step size of 0.3 degrees, and ten frame averaging. NRecon was used to binarize the 2D images before reconstructing them in the transverse plane for 3D reconstruction and volumetric analysis. Scans of bare GF and GF containing cells where set to an attenuation value allowing for imaging of the carbon substsrate. After attenuation, bare GF scans were binarized with a threshold value of 18-115 range on the contrast scale, while labelled samples were segmented into two contrast scales, one for the gold nanoparticle labelled cells (116-255), and one for the GF (18-115). 3D reconstruction of the GF environment with cells was visualized using CTVox software. Due to the threshold segmentation between the labelled cells and the GF, cells can be false colored separately allowing cellular migration and dispersion to be easily visualized with decreasing GF opacity (Figure 3A). Although a side view with the GF set to 0% opacity indicates that cells are evenly dispersed throughout the bulk of the scaffold, a top view indicates that cell density is higher towards the outer edge, which is indicative of GF’s hydrophobicity, and a characteristic we also see in fluorescence micrographs. However, fluorescence microscopy doesn’t allow for evaluations on cell migration within and on the internal structures due to the opacity of the scaffold, highlighting the benefits of utilizing the process. In addition, our CVD synthesis method for GF bioscaffolds results in microcracks in the branch sidewall as seen in SEM, leaving the internal branch structure open to cell attachment and proliferation, but difficult to confirm. Traditional methods of gold sputtering for electron microscopy and X-ray analysis would not allow for characterization of that behavior, whereas antibody staining with colloidal gold allowed us to visualize cell behavior within the branch structure (Figure 3B).

**Figure 3.**
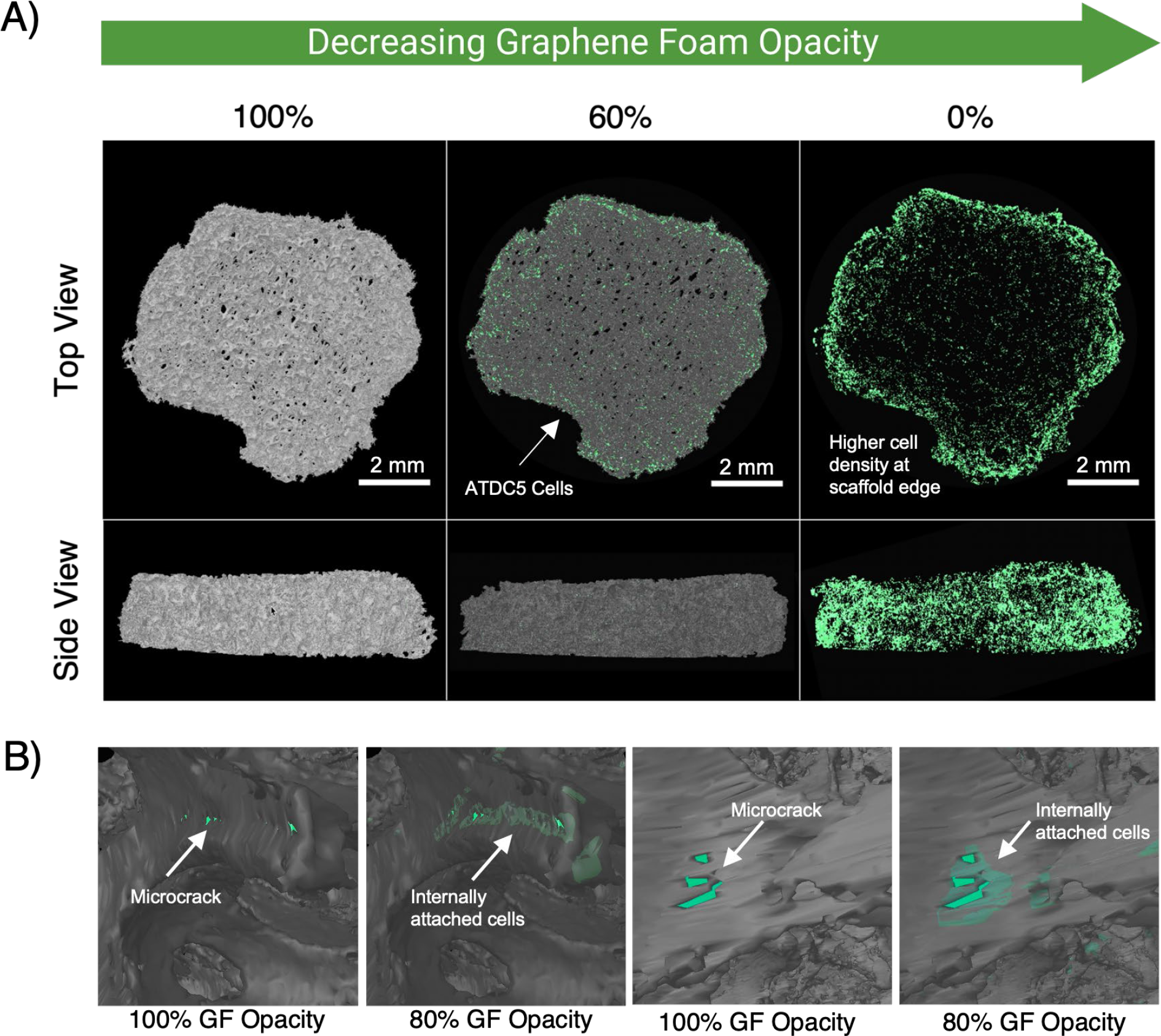
MicroCT analysis of indirectly labelled ATDC5 cells on GF. A) 3D reconstruction of GF (grey) and cells (green) showing spatial distribution in three-dimensions. B) Changing the opacity of the GF to 80% allows for visualization of internal branch cell attachment and migration through microcracks.

Indirect labelling with a conjugated fluorophore allowed us to utilize SEM, reflected light, and fluorescence microscopy techniques to characterize cell attachment and migration on the superficial plane of our GF bioscaffolds. However, it is apparent that planar analysis is not sufficient in three-dimensional environments. We have demonstrated a simple labelling technique that can be utilized for each of the aforementioned characterization modes, as well as MicroCT without any additional processing. We do not have a good understanding of the mobility of cells in optically opaque bioscaffolds like GF, and to our knowledge, a technique to asses cellular migration on GF bioscaffolds has not yet been realized. In addition, this is the first instance where cellular attachment within a branch has been confirmed. Confirmation of this activity indicates that GF bioscaffolds can be further engineered to better suit cell type and tissue organization, and through CVD we can tailor porosity and structure thickness of our resultant bioscaffolds by varying the parameters of the nickel foam template. Furthermore, the internal branch surface area of the GF bioscaffolds being available for cell attachment increases the surface area of our bioscaffolds allowing for a more robust tissue coverage. By determining how the structure of GF bioscaffolds affect cell behavior, we can design our scaffolds in such a way that we can facilitate cell organization that corresponds to tissue architecture and we can quantify how cell culture parameters effect cell behavior in 3D environments. The ability to quantify cell migration in opaque bioscaffolds is a major gap in 3D tissue engineering, as scaffolds with increased mechanical strength generally have increased opaqueness. The technique we have demonstrated is not unique to GF, and could be utilized for any bioscaffold with an optical density that can be segmented from that of gold.

## Author Contributions

D.E., J.E., and R.M-B. conceived the experiments. M.S. and O.N. fabricated the graphene foam bioscaffolds and performed cell culture, and with J.E. and J.M. performed imaging. With input from D.E., and R.M-B, M.S. and J.E. performed the data analysis. H.S. provided support for SEM imaging. M.S. wrote the manuscript with input from all authors.

## Notes

The authors declare no competing financial interest.

## Acknowledgements

This work was supported under the National Science Foundation CAREER Award #1848516. The authors acknowledge additional support from the Institutional Development Awards (IDeA) from the National Institute of General Medical Sciences of the National Institutes of Health under Grants #P20GM103408, P20GM109095, and 1C06RR020533. We also acknowledge support from The Biomolecular Research Center at Boise State with funding from the National Science Foundation, Grants #0619793 and #0923535; the M. J. Murdock Charitable Trust; Lori and Duane Stueckle, and the Idaho State Board of Education. D.E. acknowledges infrastructure support under DE-NE0008677 and joint appointment support under DOE Idaho Operations Office Contract DE-AC07-05ID14517.

